# Role of antibody-dependent enhancement in DENV-infected Wistar rats as a dengue murine model

**DOI:** 10.1101/2024.05.13.593933

**Authors:** Laura Wihanto, Cecilia Putri Tedyanto, Niluh Suwasanti, Silvia Sutandhio, Teguh Hari Sucipto

## Abstract

Preclinical studies for discovering and developing a drug for a disease involve utilizing animals as experimental subjects. The search for an effective and efficient murine model of dengue virus (DENV) infection is ongoing to support further scientific updates. This study aimed to explore the suitability of Wistar rats as a murine model for DENV infection. Twenty-four Wistar rats (male sex, 2-3 months old, 200-300 grams weight) were randomly divided into four groups (n=6 per group): control group (no infection), SC-Group (DENV-2 s.c.), IV-Group (DENV-2 i.v.), and ADE-Group (DENV-3 i.p. twice and DENV-2 i.v. once). Inactive 0.2 mL of 10^11^ FFU/mL DENV-3 were injected on days -14 and -5. Active 0.2 mL of 5 x 10^8^ FFU/mL DENV-2 were injected on day 0. Rectal temperature was measured on day 0 until 6. NS1 antigen tests were carried out from the viral medium on days -14, -5, and 0 and from the blood serum samples on day 4. Hematological parameters (leukocytes, hemoglobin, hematocrits, and platelets) were analyzed on days 0, 4, and 6. Biochemical parameters (albumin, ALT, and AST) were analyzed on day 6. SC-Group showed significant increases in the temperature from day 0 to day 1 (*p*=0.028). IV-Group showed significant increases in the temperature from day 0 to day 1 (*p*=0.007), day 2 (*p*=0.002), and day 3 (*p*=0.006). There were significant temperature increases on day 1 (*p*=0.047), day 2 (*p*=0.009), and day 3 (*p*=0.001) compared to the control group. ADE-Group had a mortality rate of 33.3%, lusterless and ruffled hair coat, and several hemorrhagic manifestations. ADE-Group also showed significant increases in the temperature from day 0 to day 2 (*p*=0.043) and day 3 (*p*=0.038). There were significant temperature increases on day 1 (*p*=0.048), day 2 (*p*=0.002), day 3 (*p*=0.000), and day 4 (*p*=0.004) compared to the control group. Leukocytes in the ADE-Group showed significant decreases from day 0 to day 6 (*p*=0.021). ALT (*p*=0.033) and AST (*p*=0.011) of the ADE-Group also showed significant increases compared to the control group. DENV infection through an induction method adapted from the antibody-dependent enhancement mechanism shows the most severe clinical manifestations and laboratory findings compared to other induction methods in Wistar rats.

## Introduction

Dengue virus (DENV) infection is a mosquito-borne disease rapidly spreading among five serotypes [1]. This positive-strand ribonucleic acid (RNA) virus from the family of *Flaviviridae* and genus *Flavivirus* is transmitted to humans through female *Aedes aegypti* and *Aedes albopictus* bites as their vectors [2,3]. The RNA viral genome encodes three structural (capsid, precursor membrane or prM, and envelope) and seven non-structural (NS) proteins, including NS1, NS2A, NS2B, NS3, NS4A, NS4B, and NS5 [4]. The NS1 is a 48-kDa multifunctional protein identified as either a mNS1 (membrane-associated or intracellular) or sNS1 (extracellular). The mNS1 plays a role in genome replication, whereas the sNS1 is secreted into extracellular and is a potential diagnostic marker for early detection of *Flavivirus* infection [4]. DENV causes a wide range of clinical manifestations in humans, manifested as asymptomatic, mild fever as in dengue fever, and even life-threatening as in dengue hemorrhagic fever or dengue shock syndrome [5,6]. The DENV-2 serotype has been reported to increase the presence and severity of plasma leakage in severe dengue, in addition to the host’s vascular integrity and the antibody-dependent enhancement (ADE) mechanism [7,8]. ADE is a process where non-neutralizing antibodies, from secondary heterologous DENV infection, will support the entry of the virus thereby increasing its binding into the host cell and its replication [8]. The incidence of DENV infection has a high prevalence worldwide, especially in tropical and subtropical regions. World Health Organization (WHO) reported that from the beginning until the end of 2023, there have been around 5,000 deaths per over 5 million DENV infection cases in more than 80 countries [9].

Providing the DENV vaccine and vector controls are preventive strategies to reduce the incidence of DENV infection [10]. Current therapy only relies on supportive treatment, with a relatively high risk of mortality if it is not sufficiently provided [11]. The current update of anti-DENV is JNJ-1802, which is in phase 2a of the human challenge model. The Janssen Pharmaceutical Companies of Johnson & Johnson is developing a safe and effective prophylactic antiviral activity against DENV-3 by blocking the NS3-NS4B interaction within the viral replication complex [12]. Antivirals for other dengue serotypes through other inhibitor pathways still need to be developed. The various clinical manifestations and complications due to severe DENV infection from different serotypes and several severity pathways of infection give rise to the urgency of searching for anti-DENV agents as specific therapies to reduce mortality rates. One of the preclinical stages in discovering and developing a drug for a disease is using animals as experimental subjects. The murine models must be able to represent conditions that mimic, are similar to, or are precisely the same as those experienced in humans. Mice (*Mus musculus*) in different strains, including AG129 (IFN α/β/γ R -/-, 9 to 12 weeks old), BALB/c (Th1/Th2, 8 weeks old), and C57BL/6 (Th1/IFN γ, 5 to 6 weeks old), are explored to be effective as murine models for experimental DENV infection [13–15]. Several limitations related to the availability of strains in some regions and the maximum blood volume produced by a mouse initiate the need to search for other more effective and efficient murine models of DENV infection.

The laboratory rat (*Rattus norvegicus*) is an animal often utilized in scientific studies due to its widespread availability, low breeding costs, and rapid metabolic rates. They produced large volumes of blood, making it more efficient to conduct research with various examination parameters. The most commonly used strain is Wistar, a sub-strain derivative of the Norway rat. These albino rats are the most popular animal model over other strains because they are widely cultivated due to their outbred history [16]. Other strains, including Sprague-Dawley, Long-Evans, and Fischer 344.

This study aimed to explore the suitability of Wistar rats as a murine model for DENV infection by examining the impact of DENV infection from several administration routes and induction methods on its clinical manifestations and laboratory findings. Several parameters are assessed, such as temperature, leukocytes, hemoglobin, hematocrits, platelets, albumin, and transaminases.

## Materials and Methods

### Ethics Statement

The animal protocols followed the ARRIVE (Animal Research: Reporting In Vivo Experiments) guidelines 2.0 and ethical principles outlined by the Council of International Organizations of Medical Sciences (CIOMS) [17]. This study was reviewed by the Health Research Ethics Committee (HREC), Faculty of Medicine, Universitas Katolik Widya Mandala Surabaya, Surabaya, Indonesia, under approval reference number 0005/WM12/KEPK/DSN/T/2024.

### DENV-2 and DENV-3 Preparation

Vero cell medium was prepared by mixing Dulbecco’s modified eagle medium (Gibco^®^, Thermo Fisher Scientific Inc., USA) and 10% fetal bovine serum (Sigma-Aldrich^®^, Merck Group, Darmstadt, Germany). Cell passage was carried out by mixing Vero cells with its medium, then centrifugated for 5 minutes using a laboratory centrifuge (Hettich^®^, Andreas Hettich GmbH & Co. KG, Tuttlingen, Germany). Centrifugation yielded two distinct layers: the media on top and the Vero cells at the bottom. The top layer was removed, and Vero cells were added to their new medium and mixed until homogeneous. A small amount of the mixture was stained with trypan blue stain 0.4% (Gibco^®^, Thermo Fisher Scientific Inc., USA) for observation and enumeration under a light microscope.

DENV-2 and DENV-3 Surabaya strains, identified by the Dengue Study Group, Institute of Tropical Diseases, Universitas Airlangga, were used in this study. A confluent monolayer of Vero cells was infected with DENV-2 at a multiplicity of infection (MOI) of 0.5 Foci-Forming Unit (FFU)/cell and then incubated at 37°C with 5% CO2 for seven days. The supernatant was collected and centrifuged for 5 minutes. The cell culture supernatant was kept at -80°C, and then the titer of DENV-2 and DENV-3 was determined using a focus-forming assay (FFA) [18,19]. The method was done multiple times until the desired titer was achieved for the dose required in the study. DENV-3 was treated by heat inactivation of 56°C for 30 minutes.

### Animal Preparation

Twenty-four male Wistar rats (2-3 months old, 200-250 grams weight) were randomly assigned to four groups (*n*=6 per group) and then divided equally into their wire-covered sterilized cage (*n*=3 per cage) with paddy husk inside as the bedding. All rats were declared healthy by the certified veterinarian and were acclimated (22±4°C room temperature, 45-55% relative humidity, 12-hour light/12-hour dark cycle) for seven days before treatment.

### Animal Experimental Design and Protocol

The experimental groups consisted of the uninfected (control group) and the infected group. Rats in the control group were not treated with any chemical intervention. Rats in the infected group consisted of three treatment groups with different virus administration routes: SC-Group (0.2 mL subcutaneous injection of 5 x 10^8^ FFU/mL activated DENV-2 on day 0), IV-Group (0.2 mL intravenous injection of 5 x 10^8^ FFU/mL activated DENV-2 on day 0), and ADE-Group (0.2 mL intraperitoneal injection of 10^11^ FFU/mL inactivated DENV-3 on day -14 and -5, then 0.2 mL intravenous injection of 5 x 10^8^ FFU/mL activated DENV-2 on day 0). The viral dosage used in this research was adapted from the previous research, with a slight of modification [20]. All experimental animals were given ad libitum access to nutritious standard chow and distilled water.

### DENV NS1 Antigen Test

A DENV NS1 antigen test was performed using a standard DENV NS1 antigen kit (StandaReagen^®^, Nantong Egens Biotechnology Co., Ltd., China). The test was carried out four times on two different samples: virus with their media and animal blood serum. The validation of DENV-3 viability was carried out on days -14 and -5, right before the virus was inactivated. The validation of DENV-2 viability was carried out on day 0, right before the injection into rats. The DENV NS1 antigen test of the animal blood serum was carried out on day 4.

### Assessment of Body Temperature

The body temperature of each rat in the experimental groups was rectally examined using a digital rectal thermometer (OMRON^®^, OMRON Corp., Japan). It was measured every day, from day 0 to 6, at the same exact time (start from the 8 a.m. daily).

### Animal Blood Collection

All rats were anesthetized with a mixture of ketamine 100 mg/kg BW and xylazine 10 mg/kg BW in a ratio of 1:1 by intramuscular injection in the caudal thigh muscle [21]. Rats were placed in the supine position and reflex observations (lack of response to a painful stimulus) were carried out first before the intervention. A 0.5 mL of blood samples were collected by cardiac puncture as the phlebotomy technique under generalized anesthesia on days 0 and 4 to perform the serial hematological analysis. On day 6, surgery was conducted to collect all the intracardiac blood samples to perform the hematological and biochemical analysis. Euthanasia was performed using the cervical dislocation technique followed by burial of the experimental animals. Animal blood collection on each of the mentioned days was carried out at the same time (after the measurement of rectal temperature), and the analyses were carried out immediately afterward without any time gap. Laboratory analyses were performed in the Veterinary Teaching Hospital (RSHP) of Universitas Airlangga, Surabaya, Indonesia.

### Hematological Parameters Analysis

Hematological parameters were analyzed on days 0, 4, and 6. Leukocytes, hemoglobin, hematocrits, and platelets were estimated using an automatic veterinary hematology analyzer (ABX Micros ESV 60^®^, HORIBA, Ltd., Japan).

### Biochemical Parameters Analysis

Biochemical parameters were analyzed on day 6. Blood serum samples were obtained from the whole blood samples following centrifugation at 3000 rpm for 5 minutes. Albumin and transaminases were estimated using an automatic clinical chemistry analyzer (Pentra C200^®^, HORIBA, Ltd., Japan).

### Statistical Analysis

All data were expressed as mean ± standard error of the mean (SEM). Statistical differences between experimental groups were analyzed using one-way analysis of variance (ANOVA) and Fisher’s least significant difference tests for the parametric analysis. Kruskal-Wallis and Mann-Whitney tests were used for the non-parametric analysis. Survival statistic was performed using Kaplan-Meier survival analysis. Statistical analysis was performed using GraphPad Prism software version 9 (GraphPad Software Inc., San Diego, California, USA). A *p*-value of less than 0.05 (*p*<0.05) indicates a statistically significant.

## Results

### Secondary DENV-infected Wistar rats presented with poorer clinical manifestations and suffered higher mortality rates compared to the control group and other DENV-infected group

No deaths occurred in Wistar rats treated with subcutaneous (SC-Group) and intravenous (IV-Group) injection routes of DENV-2. Mortality in 33.3% (two out of six) of Wistar rats was reported in the ADE-Group, in which rats were injected with DENV in different serotypes at different time intervals (Fig 1a). These rats were observed to have limping behavior and reached lethal endpoints on day 4. Wistar rat (*n*=1) representatives from each experimental group showed different clinical manifestations of their hair coats. The control group was seen in a healthy physical condition, indicated by the neat-shiny hair coat and agile behavior (Fig 1b). There were only slight changes in the SC-Group and IV-Group (Figs 1c and 1d). Lusterless and ruffled hair coat were found on the ADE-Group (Fig 1e). Differences in body posture, indicating the health status of the animal, were also observed: an upright and energetic posture (Figs 1b and 1a) compared to a limp and lethargic posture (Figs 1d and 1e). Several clinical hemorrhagic manifestations were found in ADE-Group, including hematuria (Fig 2a) and hematochezia (Fig 2c) on day 3 and ear bleeding (Fig 2b) on day 4. Chromodacryorrhea (Figs 2d1-d2, 2d4) and reddish flaky skin (Figs d2-d4) followed by minimal hair loss were also observed on day 4. These results were only found in the ADE-Group.

**Fig 1.**
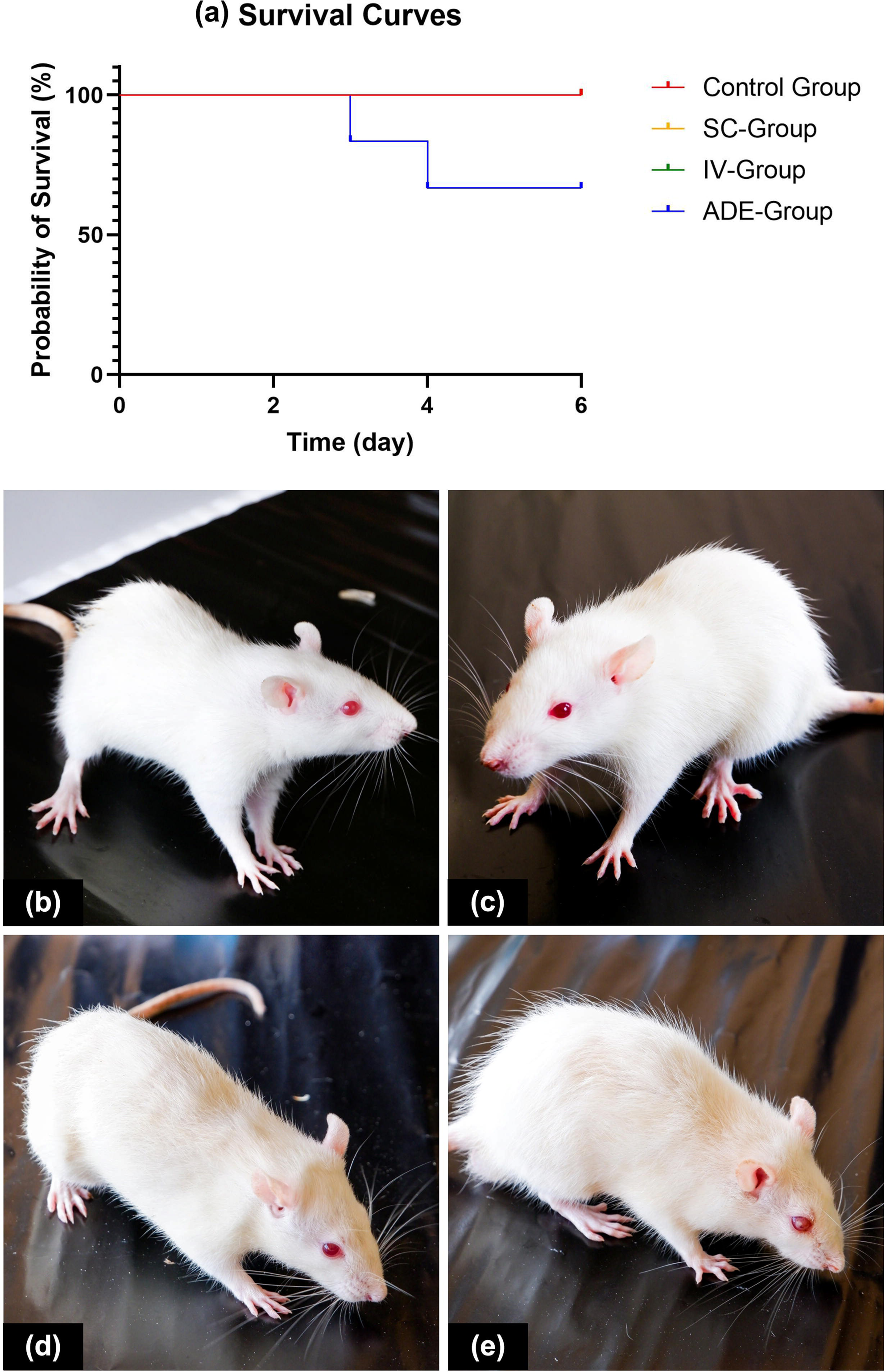
Infection of DENV in Wistar rats with different administration routes and injection methods showed a different (a) probability of survival and clinical manifestation of hair coat in (b) Control Group, (c) SC-Group, (d) IV-Group, (e) ADE-Group.

**Fig 2.**
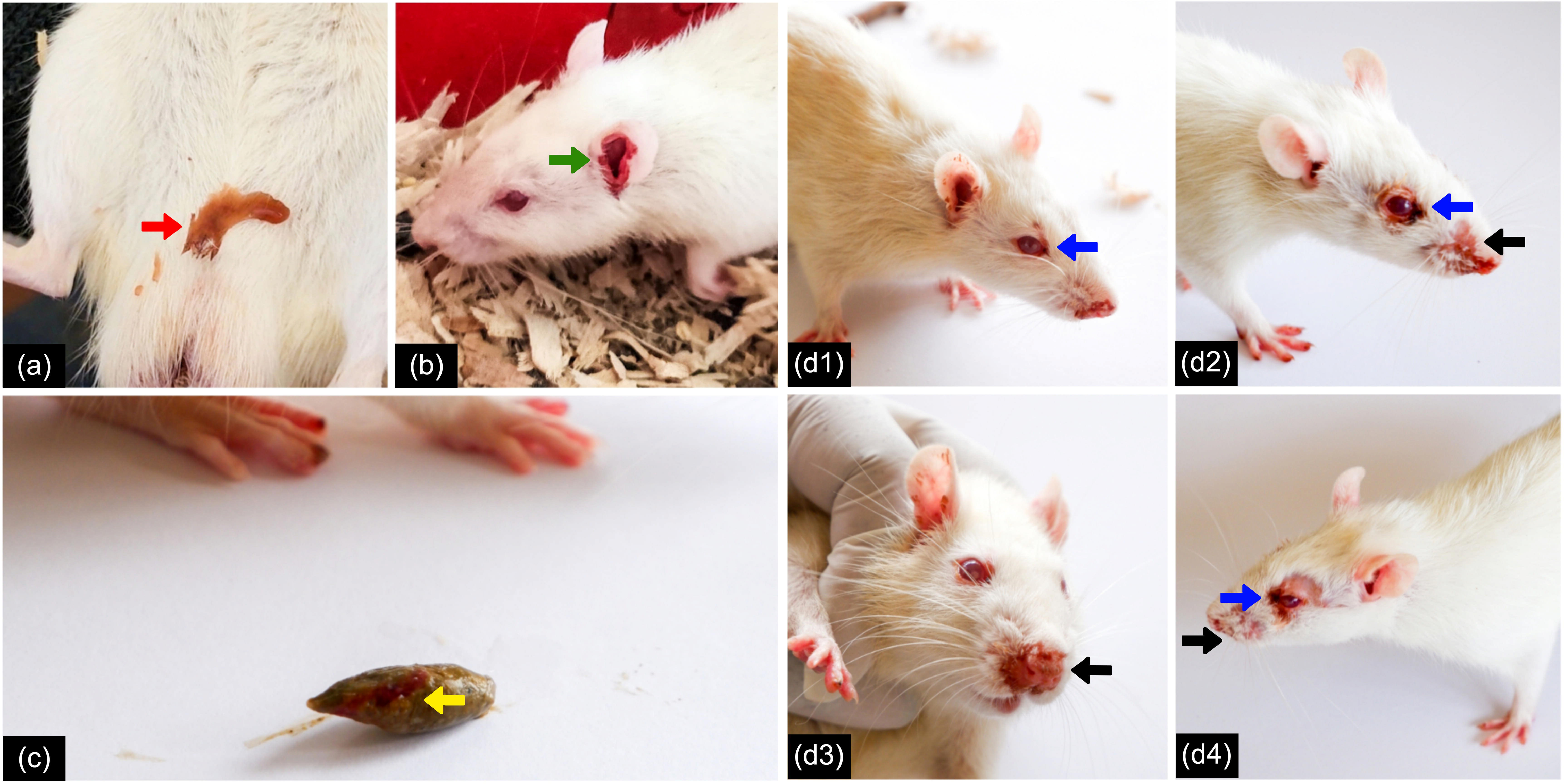
Secondary DENV-infection showed several clinical hemorrhagic manifestations in Wistar rats, including (a) hematuria (red arrow); (b) ear bleeding (green arrow); and (c) hematochezia (yellow arrow). Other findings, including (d1-2, d4) chromodacryorrhea (blue arrows) and (d2-4) reddish flaky skin with minimal hair loss around the nose (black arrows).

This result indicates that secondary DENV infection has a more severe impact on the Wistar rat than other routes of infection. Hemorrhagic manifestations in humans usually occur in the acute infection phase, similar to the hemorrhagic manifestations in Wistar rats in this study. Human bleeding tendencies could manifest as follows: purpura, petechiae, ecchymoses, hematemesis, melena, hematochezia, hematuria, or epistaxis. Thus, it can be concluded that the result also recapitulates DENV-infected human clinical manifestations.

### A conventional DENV non-structural protein 1 antigen-based test showed negative results in DENV-infected Wistar rats

Positive DENV NS1 antigen test results of DENV-3 in their medium were seen on days -14 and -5. The results were also positive on day 0, with DENV-2 in their medium as the samples. In the animal blood serum on day 4, the tests revealed negative results in all rats in the SC-Group, IV-Group, and ADE-Group (Table 1). Positive results on days -14, -5, and 0 (before injection into animals) implied that the virus used was suitable for inducing infection in this study. The negative result on day 4 in this study is inconsistent with the results in humans, while on day 4, NS1 should be positive.

**Table 1.**
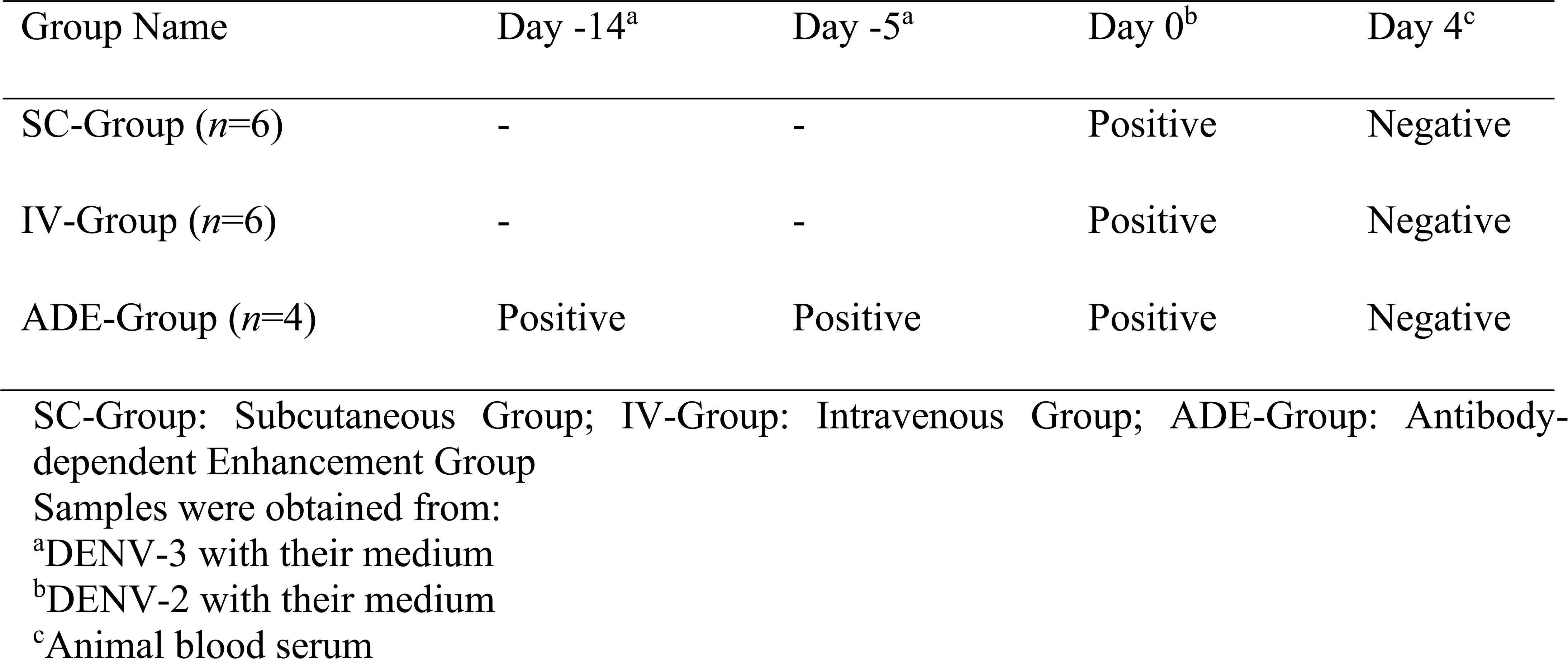
Summary of non-structural protein 1 antigen test in Wistar rat treated with DENV in subcutaneous injection (SC-Group), intravenous injection (IV-Group), and the injection method adapted from the antibody-dependent enhancement mechanism (ADE-Group)

### Acute febrile illness of secondary DENV-infected Wistar rats mimicking the fever pattern in DENV-infected human

The rectal temperature of the rats from days 0 to 6 in Fig 3 were described as follows: the control group were 35.73±0.07°C, 35.7±0.16°C, 35.78±0.17°C, 35.63±0.26°C, 35.53±0.17°C, 36±0.16°C, 35.83±0.22°C, respectively. There was no significant change from day to day (*p*=0.656); The SC-Group were 35.63±0.37°C, 36.52±0.35°C, 36±0.33°C, 35.88±0.16°C, 35.78±0.16°C, 35.82±0.23°C, 35.87±0.21°C, respectively. There was a significant increase from day 0 to 1 (*p*=0.028); The IV-Group were 35.58±0.25°C, 36.6±0.39°C, 36.8±0.14°C, 36.62±0.16°C, 36±0.22°C, 35.67±0.3°C, 35.73±0.21°C, respectively. There were significant increases on day 1 (*p*=0.007), day 2 (*p*=0.002), and day 3 (*p*=0.006) compared to day 0; and the ADE-Group were 35.68±0.5°C, 36.7±0.24°C, 37.23±0.36°C, 37.18±0.05°C, 36.68±0.42°C, 36±0.26°C, 35.53±0.84°C, respectively. Significant increases from day 0 were seen on day 2 (*p*=0.043) and day 3 (*p*=0.038).

**Fig 3.**
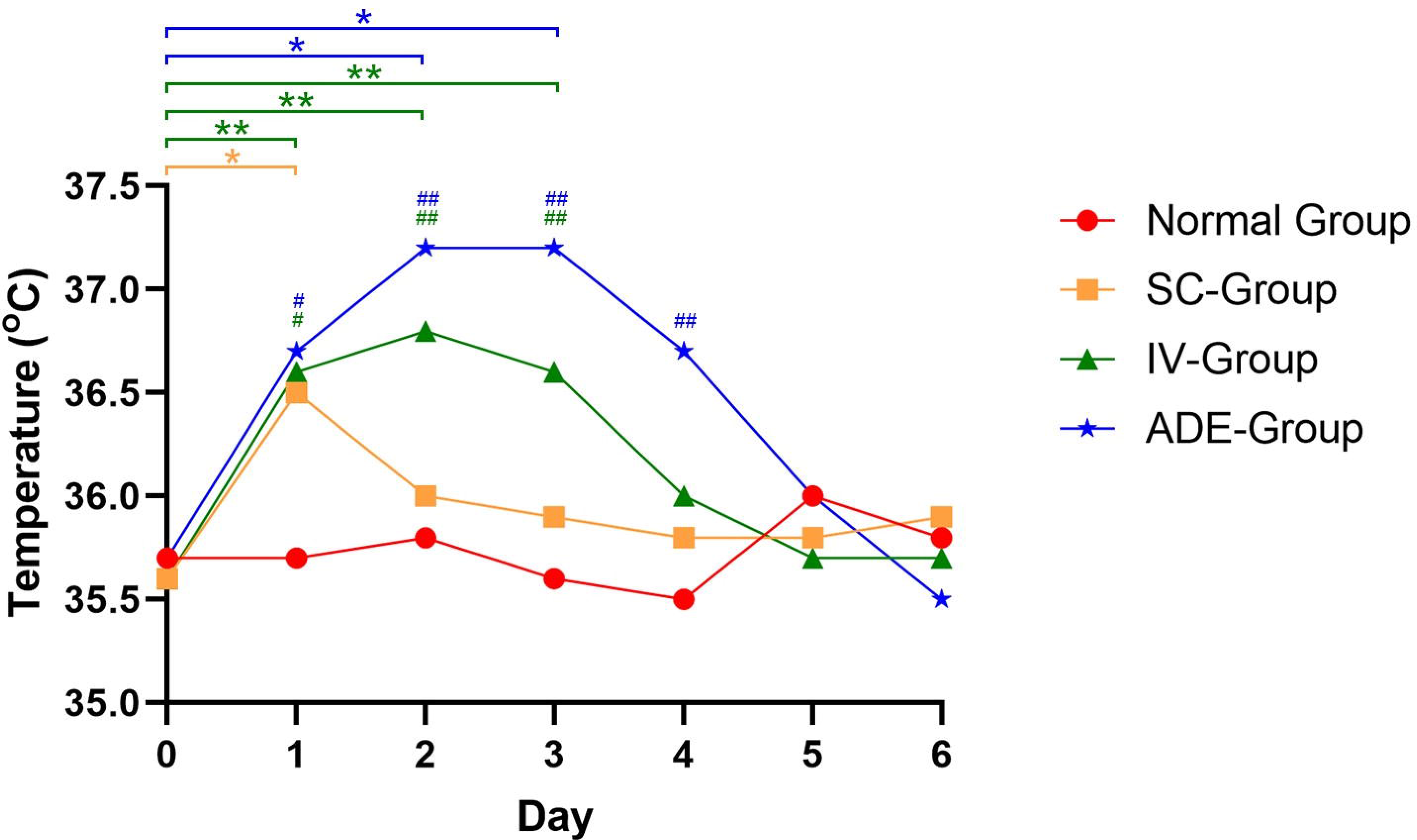
Line graph showing the rectal temperature from day 0 to day 6 of Wistar rats untreated with DENV (control group, red); treated with DENV in subcutaneous injection (SC-Group, orange), intravenous injection (IV-Group, green), and the injection method adapted from the antibody-dependent enhancement mechanism (ADE-Group, blue). *: *p*<0.05 vs day 0. #: *p*<0.05 vs control group.

If the rectal temperature in the DENV experimental group was compared with the control group, then on day 0, there was no significant difference (*p*=0.986). On day 1, there was a significant increase in the IV-Group (*p*=0.047) and ADE-Group (*p*=0.048), while in the SC-Group was not (*p*=0.069). On day 2, the IV-Group (*p*=0.009) and ADE-Group (*p*=0.002) were found to be significantly increased, while there was no significant difference in the SC-Group (*p*=0.602). On day 3, there were also significant differences in the IV-Group (*p*=0.001) and the ADE-Group (*p*=0.000) when the SC-Group still did not show a significant result (*p*=0.343). Surprisingly, on day 4, there was a significantly higher temperature that was only found in the ADE-Group (*p*=0.004) when there were no significant differences in the IV-Group (*p*=0.146) and the SC-Group (*p*=0.426). There were no significant differences between the DENV experimental group and the control group on day 5 (*p*=0.718) and day 6 (*p*=0.992).

The rectal temperature results in this study revealed that the IV-Group and ADE-Group showed a fever pattern consistent with DENV infection in humans. The fever pattern was shown to be more severe and statistically more significant in the ADE-Group than in the IV-Group.

### Secondary DENV infection decrease leukocyte counts in Wistar rats

Hematological indices of Wistar rats in all experimental groups were illustrated in Fig 4. Leukocytes (10^9^/L) in the control group were 16.9±1.59 (day 0), 16.5±1.06 (day 4), and 17.92±4.62 (day 6); The SC-Group were 11.42±0.82 (day 0), 16±1.82 (day 4), and 14.53±0.74 (day 6); The IV-Group were 17.25±2.98 (day 0), 12.77±1.09 (day 4), and 13.15±1.47 (day 6); and the ADE-Group were 23.35±1.16 (day 0), 19.7±1.68 (day 4), and 11.25±3.48 (day 6). There were no significant differences between days 0 to 4 and 6 in the control group (*p*=0.676), SC-Group (*p*=0.051), and IV-Group (*p*=0.459). In the ADE-Group, compared to day 0, was found to have no significant change on day 4 (*p*=0.083), but a significantly lower change was seen on day 6 (*p*=0.021). DENV infection in humans tends to cause leukopenia. The results of this study revealed that only ADE-Group experienced leukopenia, in which Wistar rats were injected from an injection method adapted from the ADE mechanism experienced leukopenia.

**Fig 4.**
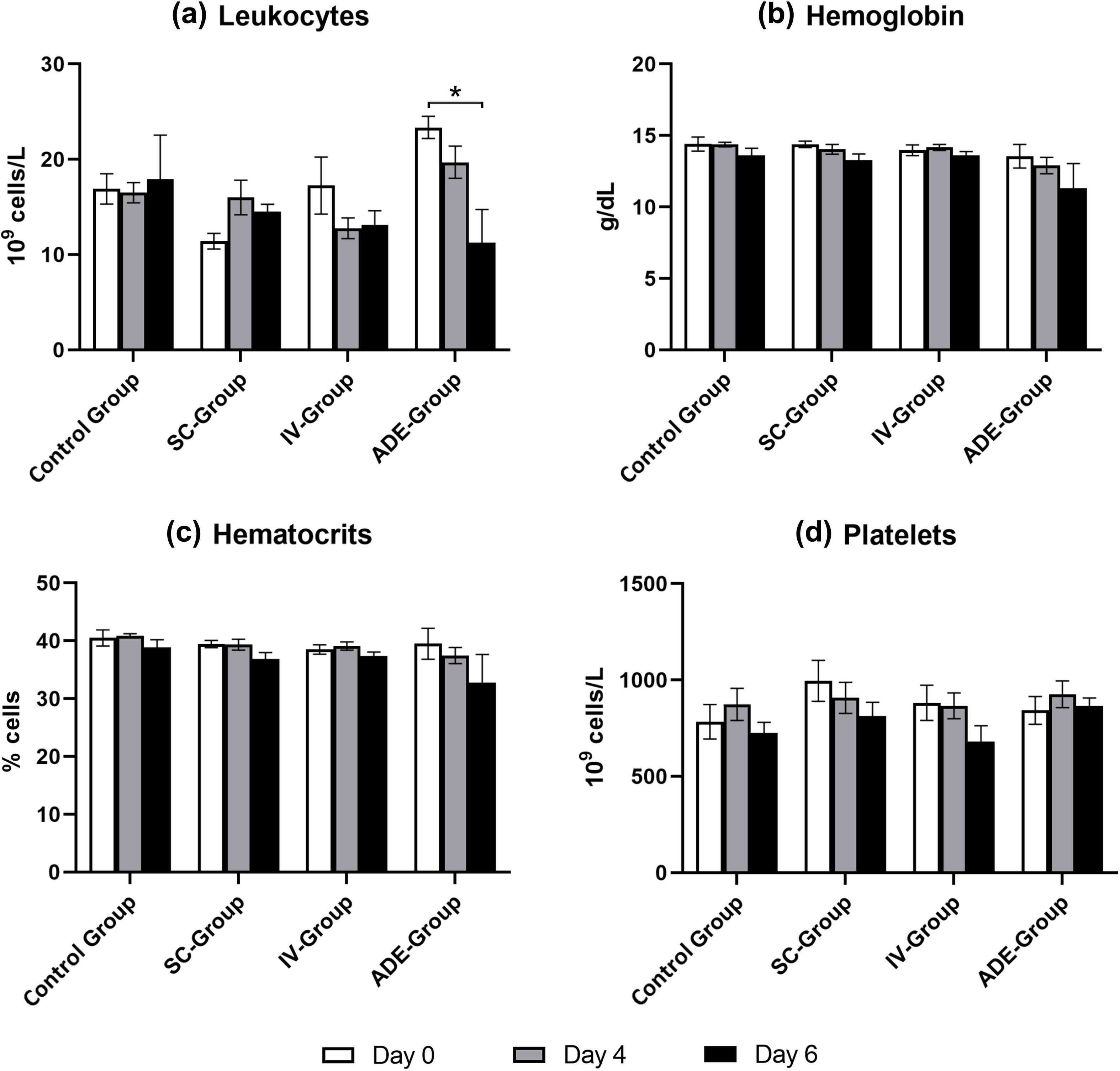
Bar diagrams of hematological parameters on day 0, day 4, and day 6, including (a) leukocytes, (b) hemoglobin, (c) hematocrits, and (d) platelets of Wistar rats untreated with DENV (control group); treated with DENV in subcutaneous injection (SC-Group), intravenous injection (IV-Group), and the injection method adapted from the antibody-dependent enhancement mechanism (ADE-Group). *: *p*<0.05 vs day 0.

Hematocrits (%) in the control group were 40.5±1.4 (day 0), 40.88±0.33 (day 4), and 38.9±1.3 (day 6); The SC-Group were 39.43±0.61 (day 0), 39.33±0.92 (day 4), and 36.88±1.11 (day 6); The IV-Group were 38.52±0.81 (day 0), 39.1±0.72 (day 4), and 37.4±0.67 (day 6); and the ADE-Group were 39.5±2.7 (day 0), 37.45±1.41 (day 4), and 32.75±4.88 (day 6). There were no significant differences between days 0 to 4 and 6 in the control group (*p*=0.433), SC-Group (*p*=0.110), IV-Group (*p*=0.281), and ADE-Group (*p*=0.378). High hematocrit is a common finding during DENV infection in humans. This study revealed that hematocrit parameters in all animal experimental groups did not experience a significant increase compared to the control group and thus did not reflect hematocrit levels to DENV infection in humans.

Platelets (10^9^/L) in the control group were 783.5±89.67 (day 0), 873.33±83.09 (day 4), and 726.5±54.46 (day 6); The SC-Group were 995.5±106.49 (day 0), 907.5±80.39 (day 4), and 812.67±71.72 (day 6); The IV-Group were 882.17±91.57 (day 0), 866.33±66.61 (day 4), and 680.67±82.16 (day 6); and the ADE-Group were 842.5±72.22 (day 0), 926.25±69.06 (day 4), and 866±40.36 (day 6). There were no significant differences between days 0 to 4 and 6 in the control group (*p*=0.421), SC-Group (*p*=0.360), IV-Group (*p*=0.180), and ADE-Group (*p*=0.633). A low platelet count or thrombocytopenia is also a common finding following DENV infection in humans. The result revealed that Wistar rats could not recapitulate the platelet counts to DENV infection in humans.

Hemoglobin (g/dL) in the control group were 14.4±0.49 (day 0), 14.37±0.16 (day 4), and 13.62±0.5 (day 6); The SC-Group were 14.38±0.23 (day 0), 14.03±0.34 (day 4), and 13.28±0.43 (day 6); The IV-Group were 13.97±0.37 (day 0), 14.17±0.2 (day 4), and 13.62±0.26 (day 6); and the ADE-Group were 13.55±0.82 (day 0), 12.9±0.58 (day 4), and 11.33±1.72 (day 6). There were no significant differences between days 0 to 4 and 6 in the control group (*p*=0.369), SC-Group (*p*=0.102), IV-Group (*p*=0.411), and ADE-Group (*p*=0.409). DENV infection in humans usually shows a high hemoglobin level. The result revealed that Wistar rats also could not recapitulate the hemoglobin levels to DENV infection in humans.

### High serum transaminases occur in secondary-DENV-infected Wistar rats

The albumin value of the control group was 3.46±0.11 g/dL, the SC-Group was 3.43±0.14 g/dL, the IV-Group was 3.5±0.13 g/dL, and the ADE-Group was 3.49±0.17 g/dL (Table 2). There was no significant result between all groups (*p*=0.984). Albumin tends to be low in DENV-infected humans. The result revealed that the albumin value of the Wistar rat does not correlate with DENV infection in humans.

**Table 2.**
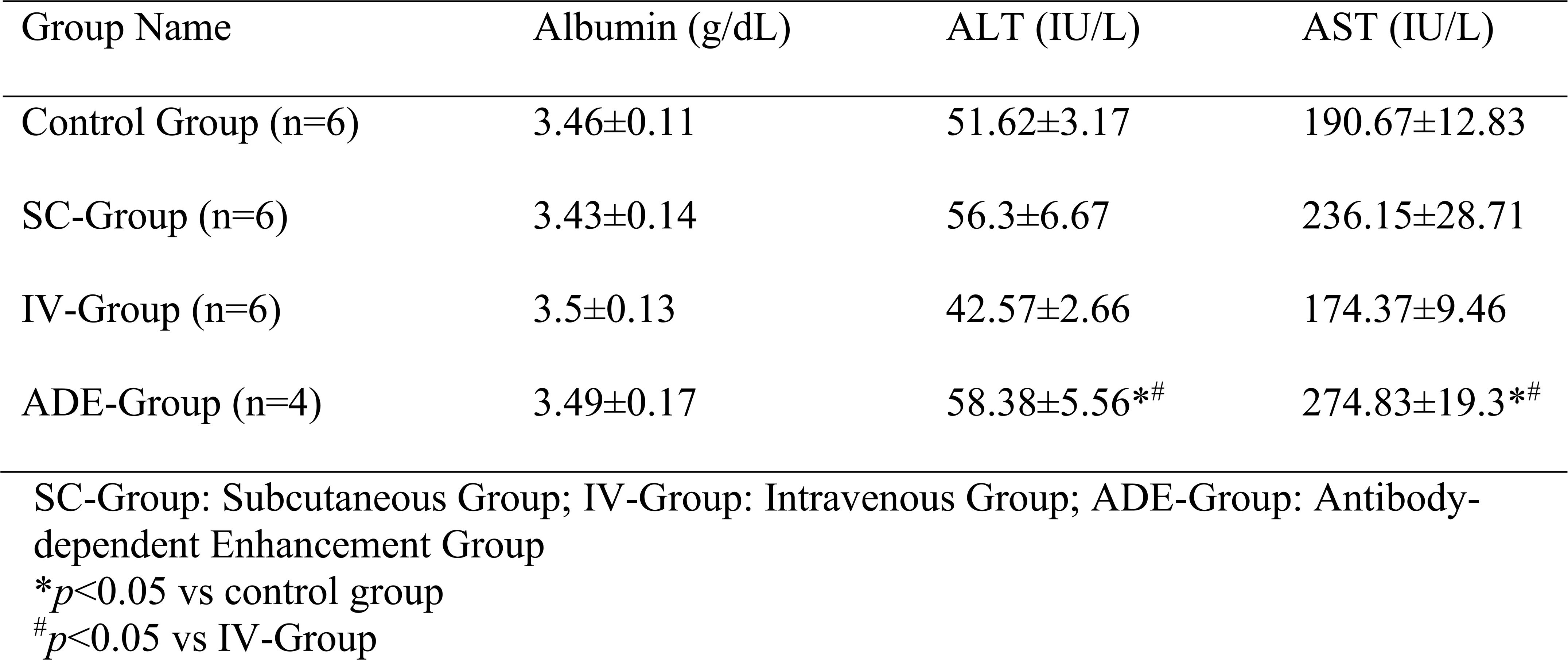
Summary of biochemical indices of Wistar rat untreated with DENV (control group); treated with DENV in subcutaneous injection (SC-Group), intravenous injection (IV-Group), and the injection method adapted from the antibody-dependent enhancement mechanism (ADE-Group)

The ALT value of the control group was 51.62±3.17 IU/L, the SC-Group was 56.3±6.67 IU/L, the IV-Group was 42.57±2.66 IU/L, and the ADE-Group was 58.38±5.56 IU/L. The ADE-Group was found to have a significantly greater ALT value than the control group (*p*=0.033) and the IV-Group (*p*=0.01). No significant differences between the SC-Group (*p*=0.749) and IV-Group (*p*=0.078) compared to the control group (Table 2). The AST value of the control group was 190.67±12.83 IU/L, the SC-Group was 236.15±28.71 IU/L, the IV-Group was 174.37±9.46 IU/L, and the ADE-Group was 274.83±19.3 IU/L. The ADE-Group was found to have a significantly greater ALT value than the control group (*p*=0.011) and the IV-Group (*p*=0.016). There were no significant differences between the SC-Group (p=0.200) and IV-Group (*p*=0.337) compared to the control group, as seen in Table 2. Serum transaminases, reflecting disease severity, were found to be similar to those of DENV infection in humans only in the ADE-Group.

## Discussion

The DENV NS1 antigen test is detectable during the acute phase, usually the first week of symptoms [22]. The examination of the DENV NS1 antigen test in Wistar rats revealed negative results in this study. As a clarification, it was confirmed that the injected virus was viable based on the positive results shown in the virus medium samples before it was injected into the rats. These results are also in line with another study, which stated that the results of the NS1 examination are not correlated with the disease severity of dengue fever in mouse model [23]. The DENV NS1 antigen test used in this study can only detect qualitative differences between negative and positive interpretations. It is possible that the NS1 antigen in the rats was at a low concentration. Thereby, this result can be deemed to be a false negative. Streptavidin-conjugated quantum dots (QDs/SA) is a highly sensitive immunosensor to detects NS1 antigen at a very low concentration (1 pM to 120 nM) [24]. The use of other methods to detect NS1 antigen in further studies needs to be considered.

There are three phases of DENV infection in humans: febrile, critical, and convalescence [25]. The febrile phase occurs on days 0 to 7, characterized by increasing temperature that reaches its peak on days 1 to 3, then gradually decreasing on days 4 to 7 [26]. This fever pattern is very similarly reflected in the ADE-Group, where the temperature rapidly increases but gradually from days 0 to 3, reaches its peak on days 2 to 3, and then gradually decreases on days 4 to 6. The fever pattern in the IV-Group also reflected the human pattern, but it was not as similar to it as in the ADE-Group. The fever pattern in the SC-Group did not reflect the DENV-infection in humans, probably because subcutaneous route infection requires an incubation period for the virus to give rise to clinical symptoms. The incubation period for DENV is 5-7 days.

The critical phase occurs on days 4 to 6 [25]. Plasma leakage, an increase in hematocrit levels of more than 20% of the basal hematocrit, might occur in this phase [26,27]. In this study, hematocrit showed no significant results in all groups. Other complete blood parameters, such as hemoglobin and platelets, did not show any significant change in all animal experimental groups. Only leukocytes showed a statistically significant decrease from day 0 to 6 in the ADE-Group, in which viral infections are usually characterized by leukopenia [28,29]. Patients are found to suffer leukopenia for several days during acute DENV infection, with an increase in lymphocyte count followed by a decrease in the absolute number of neutrophils and monocytes [30]. This was believed to be due to the destruction of myeloid progenitor cells with hypocellular bone marrow in the first seven days of the infection, in which leukocytes predominantly comprise neutrophils (around 55-70% of the total leukocyte count) [30]. The decreased neutrophil-to-lymphocyte ratio (NLR) usually happens in viral infections because monocytes and granulocytes are not essential for the clearance of a viral infection, whereas lymphocytic cells have an important role in the immune response against viruses [31].

Laboratory manifestations that can also be found in human’s DENV infection are transaminitis and hypoalbuminemia [32,33]. Increased levels of serum aminotransferase are crucial in determining the severity of DENV infection in humans [34]. It is indicated as one of the three features that characterize severe dengue, along with liver enlargement. Significant increases in ALT and AST were shown in the ADE-Group compared to the control group. ALT and AST in the ADE-Group were also found to be significant higher when compared to the IV-Group. It is implied that the secondary DENV-infection has more severity impact to the liver injury than the primary DENV-infection in Wistar rats. Surprisingly, the albumin values did not change significantly in all animal experimental groups.

Secondary DENV infection causes more severe infection reactions due to ADE [35]. The ADE mechanism has two different pathways, namely extrinsic and intrinsic [36]. Extrinsic ADE occurs due to enhanced rates of receptor interaction and internalization of virus-immune complexes, while intrinsic ADE involves the modulation of innate immune effectors by internalized virus-immune complexes to favor increased viral replication and release [37]. The intrinsic ADE also promotes the expression of pro-inflammatory cytokines, leading to increases in the production of nitric oxide, which is a radical and ultimately leads to organ damage [38,39].

The clinical manifestations, mortality rate, fever pattern, leukocytes, and transaminases were found to be the most severe in Wistar rats using the DENV infection method that utilized the ADE mechanism compared to other injection methods. The subcutaneous injection route does not provide results that are consistent with the DENV infection in humans because viruses that enter subcutaneously may require a longer research observation time, considering the DENV incubation period [40]. The intravenous injection route provides minimal insight into the disease. Further studies should be carried out in order to support these findings, such as examining the viral titers, the specific antibodies in the ADE pathway, and the pro-inflammatory cytokine or free radical parameters that also play a role in the mechanism of DENV infection.

## Conclusion

DENV infection through an induction method adapted from the antibody-dependent enhancement mechanism shows the most severe clinical manifestations and laboratory findings compared to other induction methods in Wistar rats, making it a promising DENV murine model. Future studies of possible mechanisms of infection and different parameters of other disease severity pathway are warranted to confirm our findings.

## Author Contributions

LW, CPT, NS: experimental design, conceptualization, data analysis, manuscript writing. SS, THS: methodology, research supervision, manuscript editing.

## Acknowledgements

The authors thank the Faculty of Medicine, Universitas Katolik Widya Mandala Surabaya, Surabaya, Indonesia, for their utmost support. The authors also thank the anonymous reviewers for their conscientious review of this manuscript.

